# EEG and computational aspects of how aging affects sleep slow waves

**DOI:** 10.1101/2023.12.28.573547

**Authors:** Karim El Kanbi, Núria Tort-Colet, Karim Benchenane, Alain Destexhe

## Abstract

Sleep slow-waves have been reported to vary with age in human subjects, as well as in mouse, but the underlying mechanisms remain unclear. Here, we perform a precise quantification of the effect of aging on the shape and dynamics of sleep slow waves, in a large cohort of human subjects recorded with the electro-encephalogram (EEG) during sleep. The fine-structure analysis of slow waves reveals that they slow-down, increase of variability and decrease in amplitude with age. We next investigate a computational model of the genesis of slow-wave activity and model the aging by a global decrease of the strength of the external excitatory drive to the network. This simple model reproduces some of the main features observed in the EEG, suggesting that changes of long-range excitatory connection strength may explain the evolution of slow-waves with age.

## 1. Introduction

It is commonly believed that older adults experience poorer sleep quality compared to younger individuals. However, the specific nature of this difference remains unclear. Does it manifest as reduced sleep duration, slower recovery, or increased sleep complaints? Prior research suggests that aging leads to more fragmented sleep, characterized by a decrease in the proportion of deep sleep (slow-wave sleep) and alterations in slow wave characteristics (Dijk et al., 1989, 2010; Carrier et al., 2011). However, existing studies using electroencephalography (EEG) recordings have primarily been conducted in controlled laboratory or hospital settings, limiting the generalizability of findings to real-world sleep environments. Additionally, these studies typically categorize participants into discrete age groups, hindering the examination of the continuous evolution of sleep parameters with age. This study addresses these limitations by leveraging the DREEM user database. This unique resource provides EEG sleep recordings from healthy individuals aged 19 to 85 who slept multiple nights in their natural environments. By examining individuals free from sleep disorders and controlled lab environments, this study presents a valuable opportunity to investigate the continuous evolution of sleep slow waves throughout the lifespan.

In mice, the senescence-accelerated mouse prone 8 (SAMP8) model of early aging shows disturbances in the spontaneous cortical network activity when compared to control mice (Castano Prat et al., 2017). In particular, a reduction in the frequency of the slow oscillation (SO) was found by Castaño-Prat and colleagues, together with an elongation of Up and Down states, as well as an increase in the variability of the SO cycle. Interestingly, this study also revealed a decrease in the activity levels during the Down states. We took inspiration from these experiments, to construct a computational model.

We use a previousy-introduced spiking network model of slow-wave activity (di Volo et al., 2019), where we excplicitly modeled the effect of reduced activity in Down states by reducing the external excitatory drive to the network. We examine this model in comparison to the EEG experiments.

## 2. Methods

### 2.1. EEG recordings

#### Subjects

DREEM provided a database of sleep recordings from 2377 subjects selected in the database (1678 men, 699 women), among users that agreed to participate in a research program, and 5 nights with high signal-to-noise ratio were picked up per user. Selected subjects are between 19 and 85 years old, claim to have no sleep issue, and slept with the DREEM headband in an “ecological environment”.

#### Study Device

The DREEM headband (DH) device is a wireless head-band worn during sleep that records, stores, and automatically analyses physiological data in real-time without any connection (e.g., Bluetooth, Wi-Fi, etc.). Following the recording, the DH connects to a mobile device (e.g., smartphone, tablet) via Bluetooth to transfer aggregated metrics to a dedicated mobile application and via Wi-Fi to transfer raw data to the sponsor’s servers. Five types of physiological signals are recorded via 3 types of sensors embedded in the device: 1) brain cortical activity via 5 EEG dry electrodes yielding 7 derivations (FpZ-O1, FpZ-O2, FpZ-F7, F8-F7, F7-01, F8O2, FpZ-F8; 250Hz with a 0.4-18 Hz bandpass filter); 2, 3, & 4) movements, position, and breathing frequency via a 3D accelerometer located over the head; and 5) heart rate via a red-infrared pulse oximeter located in the frontal band.

The EEG electrodes are made of high consistency silicone with soft, flexible protrusions on electrodes at the back of the head enabling them to acquire signals from the scalp through the hair. The DH is composed of foam and fabric with an elastic band behind the head making it adjustable such that it is tight enough to be secure, but loose enough to minimize discomfort. An audio system delivering sounds via bone conduction transducers is integrated with the frontal band; auditory stimulations can be delivered in real-time depending on of EEG signals. The device is described in details in (Debellemaniere et al., 2018; Arnal et al., 2020).

#### Data Analysis

##### Automatic sleep scoring

At the end of the recording, sleep data are transferred to the DREEM servers and processed automatically. The scoring of the sleep stages is done by a deep learning algorithm, which was trained on sleep recordings that were scored by several sleep experts. The deep learning architecture and the training process are described in detail in (Chambon et al., 2018).

##### Slow waves detection

Offline detection of slow waves was restricted to NREM sleep and was based on the following algorithm. This algorithm is based on a virtual channel representing one of the two fronto-occipital derivations of the sleep headband that have the best signal-to-noise ratio (Debellemaniere et al., 2018). The EEG signal of this virtual channel is then filtered in the band 0.5-18 Hz and a threshold is computed as two times the standard deviation of the filtered signal amplitude. Periods between two consecutive falling zero-crossings are detected, and only periods shorter than 300 or longer than 1400ms are discarded. Among the remaining periods, we define as slow waves those where the filtered signal crossed the negative threshold for more than 30ms.

##### Statistics

Correlation between age and sleep metrics was assessed with a Pearson correlation. Pearson’s correlation coefficient R and p-value are shown on each figure, significant correlations are highlighted.

### 2.2. Computational models

The spiking network consists of two populations of adaptive exponential integrate and fire (adex) neurons: an excitatory population of regular spiking neurons (RS) and an inhibitory population of fast spiking neurons (FS). Excitatory neurons represent 80% of the total number of cells in the network which is N = 10000. Adex neurons are described by the following equations:

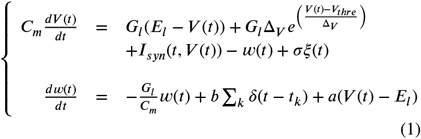

where the synaptic input *I*_*syn*_ is defined as

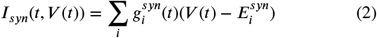

With

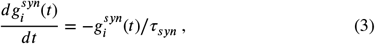

where *G*_*l*_ = 10 nS is the leak conductance and *C*_*m*_ = 150 pF is the membrane capacitance. The resting potential, *E*_*l*_, is 60 mV or 65 mV, for RS or FS cells, respectively. Similarly, the steepness of the exponential approach to threshold, V is 2.0 mV or 0.5 mV, for RS or FS cells, respectively.When V reaches the trheshold, *V*_*thre*_ = 50 mV, a spike is emitted and V is instantaneously reset and clamped to *V*_*reset*_ = 65 mV during a refractory period of *T*_*refrac*_ = 5 ms. Every neuron receives an identically distributed white noise *ξ*(t) of zero mean and instantaneously decaying autocorrelation. The noise amplitude is 50. The membrane potential of RS neurons is also affected by the adaptation variable, w, with time constant *τ*_*w*_ = 500 ms, and the dynamics of adaptation is given by parameter a = 4 nS. At each spike, w is incremented by a value b, which regulates the strength of adaptation. In order to simulate a strong adaptation, which aims at modelling the effects of deep anesthesia, the parameter b is set to 60 pA in RS cells, while FS have no adaptation. Excitatory and inhibitory conductances are 1 and 5 nS, respectively, with a time constant *τ*_*syn*_ = 5 ms and reversal potentials *E*_*syn*_ = 0 mV or 80 mV for excitatory and inhibitory synapses, respectively. The external drive that we use to bring our model into the control or aged condition targets all neurons in each population and consists of a poissonian source of 0.15 Hz or 0.05 Hz in the control or the early aging condition, respectively. Each simulation was 200 s long, and had a time step of 0.1 ms.

## 3. Results

We start by showing the results in the human EEG, then we show the computational model.

### 3.1. Human EEG

#### Changes in sleep architecture: sleep tends to be more fragmented with age

It has been reported that older people have more wake episodes between sleep onset and awakening in the morning and thus the sleep efficiency is also affected (Dijk et al., 2010; Landolt and Borbély, 2001). Here in their ecological environment, subjects choose the time to go to sleep and then we measure their usual sleep metrics (Fig. 1).

**Figure 1:**
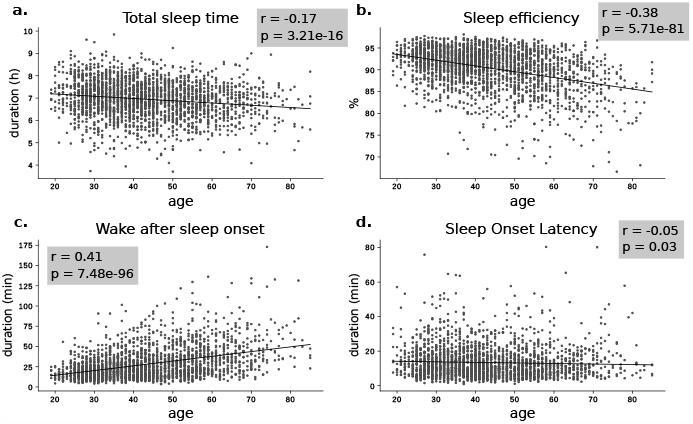
Evolution of sleep standard metrics: sleep fragmentation increases with age: (a) Total sleep time (TST) (b) Sleep efficiency is the percentage of TST over Time in Bed (TIB) (c) Wake after sleep onset (WASO) is the total duration of wake episodes between sleep onset and the awakening (d) Sleep Onset Latency (SOL) is the duration before the first consolidated sleep episode.

We also observe a significant increase in Wake after sleep onset (WASO), defined as the duration of wake periods after sleep onset, as well as Total Sleep Time (TST) and Sleep Onset Latency (SOL), although the effect of ageing on SOL is not obvious. There is a clear effect of ageing on sleep efficiency, as we observe a decrease with a correlation coefficient of -0.38.

#### Age-Related Decline in N3 Sleep with Subtle Decreases in REM Proportion

We observed a pronounced and significant reduction in the proportion of N3 sleep, accompanied by a marginal decrease in REM sleep proportion. This shift explains the observed increase in N2 and N1 stages (Fig. 2). While the overall balance between NREM and REM sleep appears to remain constant throughout life, a notable decrease in Slow Wave Sleep within the NREM stage is evident. This significant reduction of slow waves in older individuals corroborates findings from various studies (Dijk et al., 1989; Landolt and Borbély, 2001; Ohayon et al., 2004; Van Cauter et al., 2000; Mander et al., 2017). However, the change in REM sleep has been less clear, with some studies indicating a slight decrease and others reporting no significant change.

**Figure 2:**
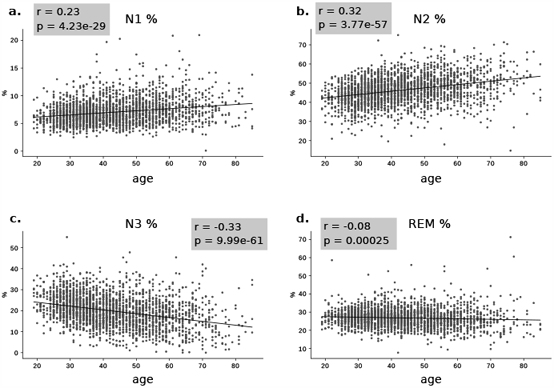
Strong decrease of slow-wave sleep proportion. Evolution with age of the percentage of each sleep stage: (a) N1, (b) N2 (c) N3, (d) REM. No significant change is observed in the percentage of REM sleep among the subjects, but we observe a decrease of N3 proportion in favour of N1 and N2 stages.

Our analysis also shows an increase in wake episodes during sleep and a decrease in NREM sleep depth. As sleep stages offer a macroscopic view of NREM sleep dynamics, it’s essential to analyze the evolution of its primary cortical rhythms, slow waves and spindles, to gain a deeper understanding of the mechanisms underlying these alterations.

#### Slow waves become flatter for older subjects

To assess the evolution of sleep slow waves from young to older subjects, we first quantified four shape characteristics often described in sleep studies. These four measures: negative and positive amplitudes, and duration (or width) could be correlated to homeostasis processes and down states characteristics (McKillop et al., 2018; Riedner et al., 2007). As studies described a decrease in slow wave amplitude with age (Carrier et al., 2011), our slow wave detection method was not based on a unique threshold in millivolt, instead, the threshold was a factor of the standard deviation of the EEG signals. Here in Fig. 3, all slow waves features show a flattening of slow waves with age, which could be interpreted as a decrease in down states amplitudes, or a sparseness of down states occurrences. While we have no access to neuronal information with the EEG, we can quantify the evolution of the local density of slow waves in our subjects.

**Figure 3:**
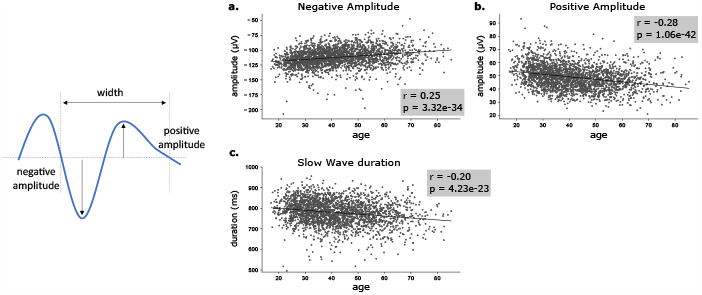
Flattening of slow waves with aging. Left Slow Waves characteristics: amplitudes are expressed in mV and durations in milliseconds. Right Evolution of slow waves characteristics averaged per subjects (a) Negative amplitude of slow waves. (b) Positive amplitude of slow waves. (c) duration of slow waves which is the width between the two consecutive zero-crossing. (d) rising slope of slow waves after the trough.

#### The occurrence of slow waves is denser and more regular for young subjects

A simple way of quantifying the density of slow waves in a sleep record, one can simply look at the number of slow waves divided by the time in sleep or NREM, as shown in Fig. 4*a*, where we observe a decrease in both the number of slow waves and their density with age. However, one can have a better sight of their occurrences by quantifying the Inter Slow-waves Intervals, defined as the distance between two consecutive slow waves; three examples of ISI distributions are shown in Fig. 4*b*, for single nights of three subjects of 27, 36 and 52-year-old. In these three examples, we note that the distribution is tighter for the younger subject and that the older subject has a bigger peak value. It is quantified for all subjects in Fig. 4*c*: the peak of the ISI distributions increase with age, thus slow waves are sparser as we become older.

**Figure 4:**
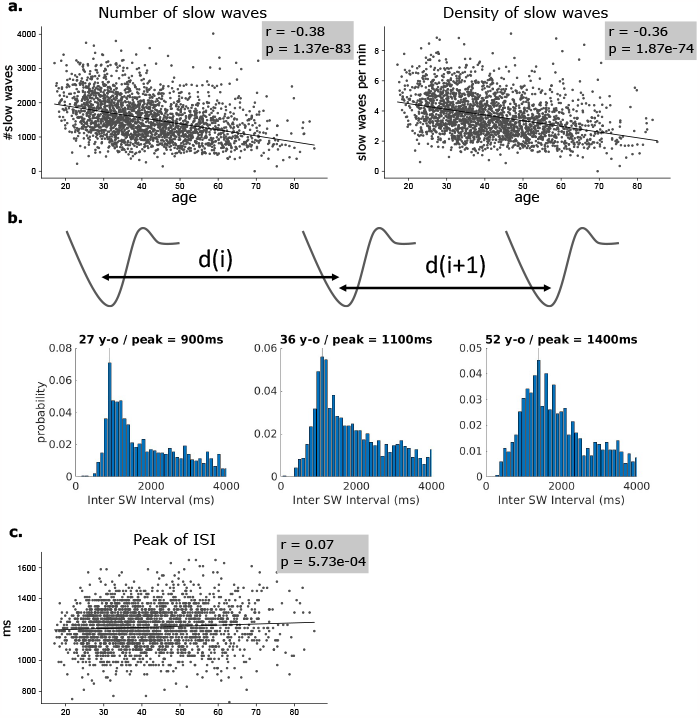
The occurrence of slow waves during sleep is sparser and less regular with age. (a) Evolution of the number and the density of slow waves with age, averaged per user (b) Schematic representation of the inter slow-waves intervals (ISI) quantified. Example of three distributions of ISI for three sleep recordings (27, 36 and 52 y-o). (c) Evolution with age of the mode (peak) the ISI distribution: the increases with age show that slow waves tend to be less dense and less regular.

#### Strong evolution of slow waves homeostasis is due to a difference at the beginning of the night

Several studies reported a difference in sleep propensity between young and older subjects, maybe reflecting a lessening in homeostatic requirement. To compare these observations with our data and add a new approach, we measured for each night the evolution of slow wave occurrence (per minute) and compute slow wave homeostasis curves (Fig. 5*a*: examples of two homeostasis curves for young and old subjects). The slope of this curve gives us an indication of the homeostatic process S, which is supposed to have a higher amplitude in young subjects (Landolt and Borbély, 2001; Mander et al., 2017; Landolt et al., 1996). The decrease of the slope value, observed across the lifespan, is coherent with the literature, also in our continuous age-related dataset, we do not seem to observe a breaking point where this homeostatic change would appear (Fig. 5*b*). Then a question remains: is this difference in the homeostatic requirement due to a change at a specific moment of the night? Indeed, we see a strong effect of ageing on the number of slow waves in the first two hours of sleep, and small but significant effect concerning the last two hours (Fig. 5*c*). Therefore, if an external enhancement of slow waves were to be done to improve sleep in older people, it should be tried in the first hours of sleep.

**Figure 5:**
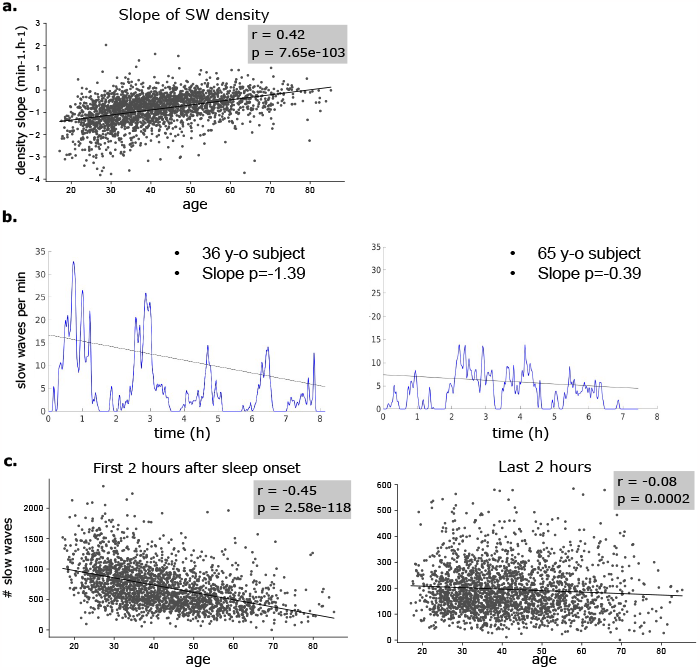
Slow waves homeostasis changes occur in the first hours of sleep. (a) Example of slow waves homeostasis curves for two sleep recordings (left young subject, right young subject), slopes of this density curves are measures of sleep homeostasis. (b) Evolution of the slopes of slow waves homeostasis curves with age. (c) Significative strong effect of ageing on the proportion of slow waves is observed in the first two hours of sleep, yet smaller effect in the last two hours. The difference in homeostasis slope in b is explained by a strong decrease at the beginning of the night.

#### Effect of the auditory stimulations of N3

In addition to recording physiological data during sleep, DREEM device has been designed to deliver auditory stimulations in N3 to induce slow waves. A subset of subjects (n=300, 200m) received around 50% of auditory stimulations and 50% of sham (i.e., stimulation with no sound), stimulations are triggered in N3 right after the detection of a slow wave. The process is described in detail in Debellemaniere et al. (2018). First, we looked at the instantaneous effect of auditory stimulations on EEG (Fig. 6*a*): auditory Evoked Related Potentials of stimulations is significative bigger than for sham. To assess the effect on slow waves induction of sounds, we assessed the probability of a sound to be followed by a slow wave: in the first 500ms after sound delivering, the probability of observing a slow wave is higher than chance, which is represented by sham distribution (Fig. 6*b*). This allows us to define a criterion for qualifying a stimulation of successful, it must be followed by a delta in the following 500ms.

**Figure 6:**
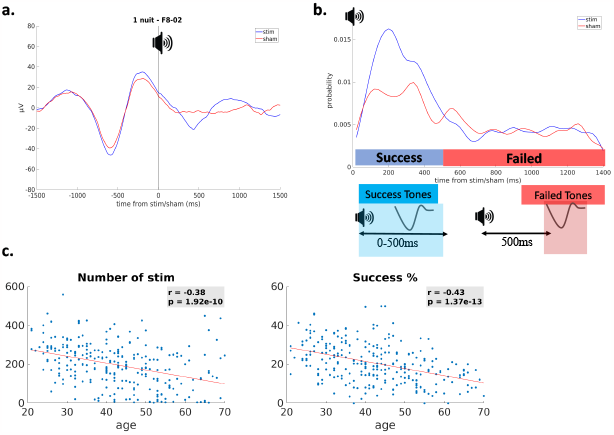
Closed-loop auditory stimulations during N3. (a) Average ERP time-locked to auditory stimulations and sham. Stimulations and sham are delivered just after the detection of a slow wave. (b) Distribution of the delay between auditory stimulations or sham and the first following slow wave. Stimulations increase the probability of slow wave occurrence in the first 500ms: successful stimulations are defined as tones followed by a slow wave within 500ms. (c) Number of stimulations per sleep recording and percentage of successful stimulations.

As older subjects have less slow waves, the closed-loop stimulation process delivers fewer stimulations for older subjects and more for young subjects; however, the percentage of successful also strongly decreases across the lifespan (6*c*). Thereby, as age increases, the endogenous and the external induction of slow waves decreases, maybe reflecting a change in the cortical network that would lead to less down states induction.

### 3.2. Computational model

To model these aging effects on slow-wave activity, we simulated a spiking network model introduced previously (di Volo et al., 2019), which consists of a network of excitatory (FS) and inhibitory (FS) neurons described by the Adaptive Exponential (AdEx) integrate and fire model (see Methods). Experimental observations in SAMP8 mice Castano-Prat et al. (2017) revealed that aging is associated with a decrease of spontaneous activity in the Down states of slow-waves. In our model, the spontaneous activity is provided an external excitatory drive to the network (see Methods), so we modeled aging as an overall reduction of this excitatory drive.

Accordingly, the network model displayed a reduced spontaneous activity when decreasing the excitatory drive (from 0.15 Hz to 0.05 Hz) targeting both populations of neurons. This reduction in the network activity during the Down states lead to an overall reduction of the network excitability, such that the frequency of the SO was reduced by 32.4% (from 1.11 ± 0.08 Hz in the control condition to 0.75 ± 0.07 Hz in the aged condition) (Fig. 7). A quantification of the slow waves in the model (Fig. 8) shows that Down state duration suffered an increase of 61.3% (from 689.7 ± 271.8 ms in the control condition to 1112.7 ± 376.3 ms in the aged condition) since the ability of the network to trigger Up states was affected by the lack of excitability. At the same time, although to a lesser extent, Up states were also elongated. The relative change was of 5.8 % (from 94.3 ± 38.9 ms in the control condition to 99.8 ± 34.7 ms in the aged condition). Finally, we observed a 28.6 % increase in the variability of the SO, computed as the coefficient of variation of the length of the Up-Down state cycle (from 0.07 in the control ondition to 0.09 in the aged condition).

**Figure 7:**
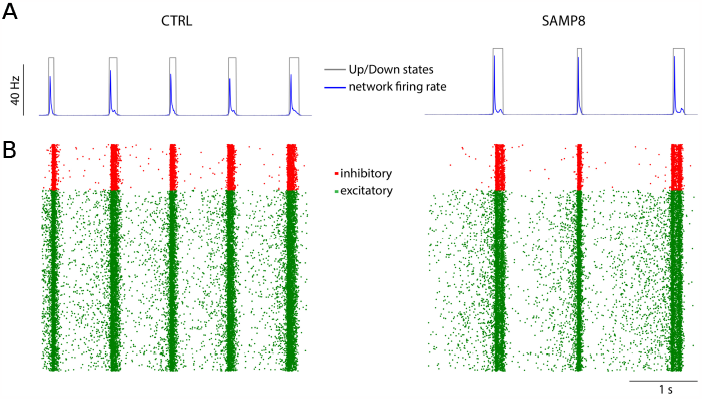
Computational model of the effect of aging on slow waves. **A**. Firing rate of the AdEx network and detection of Up and Down states (blue and gray traces, respectively) during 4s of the simulation in the control (left) and the aged (right) conditions. **B**. Raster plot of the spikes of inhibitory and excitatory cells (red and green dots, respectively) during the same 4s of the simulation as in **A** in the control (left) and the aged (right) conditions.

**Figure 8:**
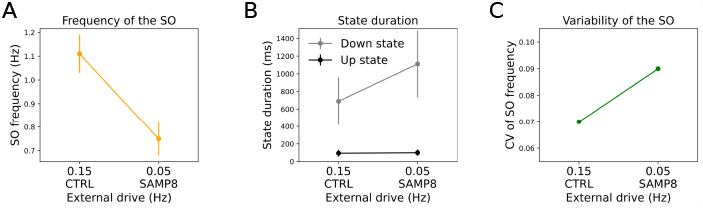
Quantification of the slow waves generated by the model. **A**. Average SO frequency as a function of the external drive. **B**. Average Up and Down state duration as a function of the external drive. **C**. CV of the SO frequency as a function of the external drive. Error bars represent standard deviation. SO: slow oscillation; CV: coefficient of variation.

## 4. Discussion

In our study, we have conducted a detailed analysis of EEG recordings during sleep from a large cohort of human subjects. This analysis has brought to light significant agerelated changes in sleep slow waves, including a slowdown in frequency, a reduction in amplitude, and an increase in variability. Alongside this empirical analysis, we also introduced a simple computational model where these features could be reproduced. We discuss below these findings.

### 4.1. Analysis of EEG recordings of slow-waves at different ages

Decades of intensive research have illuminated the impact of aging on sleep duration, architecture, and specific oscillations. Most of these studies have been conducted with subjects or patients sleeping in controlled environments, such as labs or hospitals, using uncomfortable polysomnographic systems. Our study stands out by utilizing neurophysiological data from individuals sleeping in their natural environments over several nights, with a consistent polysomnographic system, and spanning a continuous age range from 20 to 70 years old. Our findings corroborate many previously observed trends, such as increased sleep fragmentation, a shift from SWS to lighter sleep stages (N1 and N2), a reduction in homeostatic sleep pressure, and the attenuation of slow-wave activity. We also uncovered less obvious aspects of sleep and aging. The literature has been inconclusive regarding the evolution of REM sleep; however, our analysis indicates a subtle but significant effect of aging on REM sleep percentage. Additionally, the efficacy of auditory stimulation in enhancing slow waves appears to diminish with age.

A notable observation from our study is the absence of a distinct ‘breaking point’ in the progression of sleep parameter changes with age. This raises questions about whether the observed decrease in slow wave activity and increase in sleep fragmentation are gradual or if they manifest abruptly at a certain age, potentially varying among individuals. Longitudinal follow-up with our subjects over several years will be necessary to address this.

Moreover, our study highlights a concurrent decrease in slow wave homeostatic drive (particularly at the start of the night) and a reduced probability of slow wave induction through auditory stimulation. Future research could explore strategies to counteract the decline in slow waves in older individuals, potentially through auditory stimulations. However, these findings may suggest an age-related alteration in cortical network connectivity, affecting both the natural and induced generation of down states, a neuronal phenomenon integral to slow waves.

### 4.2. Computational model of the effect of aging on slow-waves

In agreement with our findings in the EEG from human sleep, our computational model shows that, as we age, the slow waves become less prominent, more variable and the Down states (which could be interpreted as the inter SW intervals) become longer, as observed here in the human EEG and previously in a mouse model of early aging Castano-Prat et al. (2017). However, a word of caution is needed here about the comparison with mice, since the mice experiments were done under anesthesia while the EEG recordings that we report here were done in natural sleep. There is evidence that the slow waves of sleep are different than those under anesthesia (Nghiem et al., 2020). Nevertheless, our model is based on a reduction of the spontaneous activity observed with aging in mice, and does reproduce some of the main features of how aging affects slow waves, such as the slowdown, the decreased synchrony and increase of variability.

Concerning the possible mechanisms, we investigated here one mechanism that accounts for the reduced spontaneous activity, based on a reduction of the external excitatory drive. It must be noted that the network model we propose here represents a local population of neurons, while the EEG is associated with large-scale effects. The excitatory drive represents the excitatory interactions provided by other cortical regions onto the local network under study, so a reduction of this drive would mean a reduced long-range excitation in the system. This constitutes the main prediction of the model, which could be tested experimentally or by large-scale models of cortical slow-wave activity (Goldman et al., 2022).

## Acknowledgments

Research supported by the CNRS and the European Union (Human Brain Project H2020-785907, H2020-945539).

## Code Availability

The code used for the simulations presented in this paper is available under request and will be available online after publication.

